# BayesCurveFit: Enhancing Curve Fitting for Early-Stage Compound Screening Using Bayesian Inference

**DOI:** 10.1101/2025.02.23.639769

**Authors:** Niu Du, Yang Chen, Yu Qian

## Abstract

Accurate curve fitting is central to quantifying dose–response relationships in drug discovery. However, commonly used regression-based methods are ill-suited for early prescreening assays, where measurements are sparse and experimental noise can dominate. Bayesian approaches have been developed previously, but they are rarely optimized for such data-limited, noise-prone conditions. We present *BayesCurveFit*, a Bayesian framework specifically designed for robust dose–response inference under data scarcity. The workflow integrates calibrated initialization, stochastic optimization, adaptive posterior sampling, and probabilistic mixture modeling within a unified Bayesian pipeline, enabling reliable parameter estimation and uncertainty quantification from few observations. By modeling residuals with a Gaussian–Laplace hybrid distribution, *BayesCurveFit* remains robust to outliers where ordinary least squares and conventional regression methods fail. Simulation studies and real screening benchmarks demonstrate that *BayesCurveFit* outperforms state-of-the-art regression-based methods in recovering true dose–response relationships under limited sampling. In addition, a Bayesian measure of significance – posterior error probability – offers interpretable probabilistic confidence for response classification. Together, these features establish a general and easy-to-use Bayesian framework for analyzing dose–response data in early screening experiments.

## 1 Introduction

Compound screening for drug discovery prior to preclinical development involves a series of filtering stages, including high-throughput screening (HTS), hit to lead (H2L), and lead optimization (LO). The number of compounds tested decreases dramatically across these stages, from 10^4^ to 10^6^ in HTS to only 10 to a few hundred compounds in LO. Due to budgetary and logistical constraints, the number of experiments conducted in the early stages of screening is necessarily limited. HTS is typically performed at a single concentration, and hit confirmation assays use only a few doses. Unlike late-stage screening, early-stage screening emphasizes sensitivity or minimizes false negatives to ensure that potentially active compounds are not prematurely excluded, whereas late-stage screening focuses on reducing false positives to identify truly effective candidates through more extensive experimentation. In most drug discovery pipelines, analytical and regulatory efforts concentrate on mature lead candidates that have advanced to preclinical or clinical evaluation. As a result, regulatory guidance and validation frameworks are primarily designed for late-stage assays and datasets supporting those downstream decisions [European Medicines Agency, 2011, U.S. Food and Drug Administration, 2018]. However, the early stage decision has consequences that are asymmetric in downstream discovery. Failing to advance a viable lead candidate during early screening carries large downstream opportunity costs for R&D programs. Analyses of drug development costs indicate that false negatives at this stage can translate into millions, even billions, of dollars in lost value [DiMasi et al., 2016, Wouters et al., 2020].

Early-stage screening usually operates under strict throughput and resource constraints, prioritizing efficiency and cost-effectiveness over replication depth and concentration coverage. In practice, the layout of microtiter plates and the use of multichannel pipetting limit the number of concentration levels that can be tested for each compound or target during early screening [Inglese et al., 2007]. Consequently, dose–response curves in early screening are typically constructed from only a small number of tested concentrations, often a few to a dozen, and without replicates [Sebaugh, 2011, Holland-Letz and Kopp-Schneider, 2015]. Such data-limited assay designs may be sufficient for ranking compound potency but brings in uncertainty in parameter estimation for curve fitting, making key metrics such as *IC*_50_*/EC*_50_ values, Hill slope, and response range highly sensitive to individual measurements.

The limited number of observations and requirements on high sensitivity in early screening experiments expose a niche largely overlooked by existing curve-fitting methodologies, which primarily rely on regression. Despite major advances in nonlinear regression modeling, few of them are designed to address the unique conditions of early screening: limited data points, heteroscedastic noise, and the presence of occasional outliers. Bayesian statistics provide a rigorous alternative for addressing these challenges by incorporating prior knowledge and providing a probabilistic interpretation of model parameters. Rather than minimizing the mean squared error as in regression analysis, a Bayesian approach known as maximum a posteriori (MAP) maximizes the posterior probability of the parameters based on the given data (likelihood), which usually excels in situations with limited data or outliers, as it supports more reliable model selection and parameter optimization [van de Schoot et al., 2021, Kruschke and Liddell, 2018].

However, existing Bayesian tools are either not specific to dose-response data analysis or not focused on early prescreening. General Bayesian libraries such as *PyMC* and *Stan* [Salvatier et al., 2016, Carpenter et al., 2017] provide flexible probabilistic building blocks but are neither fine-tuned nor assembled into turnkey solutions for dose–response inference. Domain-specific Bayesian frameworks have been developed in pharmacometrics and toxicology, but focusing on well-controlled experimental designs typical of late-stage or confirmatory studies [Pinheiro et al., 2014, Gould, 2019]. Benchmark-dose systems such as BBMD [Shao and Shapiro, 2018, Ji et al., 2022] and its genomic and omics extensions [Zilber et al., 2025] quantify uncertainty in population-level endpoints, while hierarchical dose–response meta-analyses pool replicated data to infer group-level effects [Allen et al., 2020, Hamza et al., 2021]. Even specialized Bayesian analyses for high-throughput drug screens emphasize cross-plate or cross-cell-line information sharing [Shterev et al., 2018, Labelle et al., 2019, Tansey et al., 2022], rather than for improving robustness of single-curve inference under limited sampling. More recently, learning-based approaches such as neural networks have been proposed for dose–response prediction [Campana et al., 2024]. All these methods emphasize predictive accuracy in data-rich settings and generally lack the explicit probabilistic inference and calibrated uncertainty quantification that underpin Bayesian estimation.

## 2 Methods

The design of *BayesCurveFit* is motivated by two statistical and experimental realities in early screening. First, classical asymptotic results that justify Gaussian-based inference require moderate-to-large numbers of independent measurements; with very small sample sizes, sampling distributions can deviate substantially from normality, and parameter estimates can become biased or highly variable [Kwak and Kim, 2017, Smid et al., 2020]. Second, low-frequency but real operational errors, from pipetting imprecision and occasional air bubbles to transient environmental fluctuations, introduce asymmetric, heavy-tailed, or mixed-distributed noise that violates standard Gaussian error assumptions and increases the likelihood that a promising compound’s curve will be poorly fitted [Guan et al., 2023]. To address these two challenges, we carefully selected Bayesian statistical modules and assembled them into a unified pipeline, before optimizing the pipeline for dose–response curve fitting. The pipeline integrates stochastic optimization, adaptive posterior sampling, and probabilistic mixture modeling for optimal inference (**Figure. 1**). Below we summarize the main steps and their roles, while mathematical expressions and usage guidelines are provided in **Supplementary Information (SI)**.

**Figure 1.**
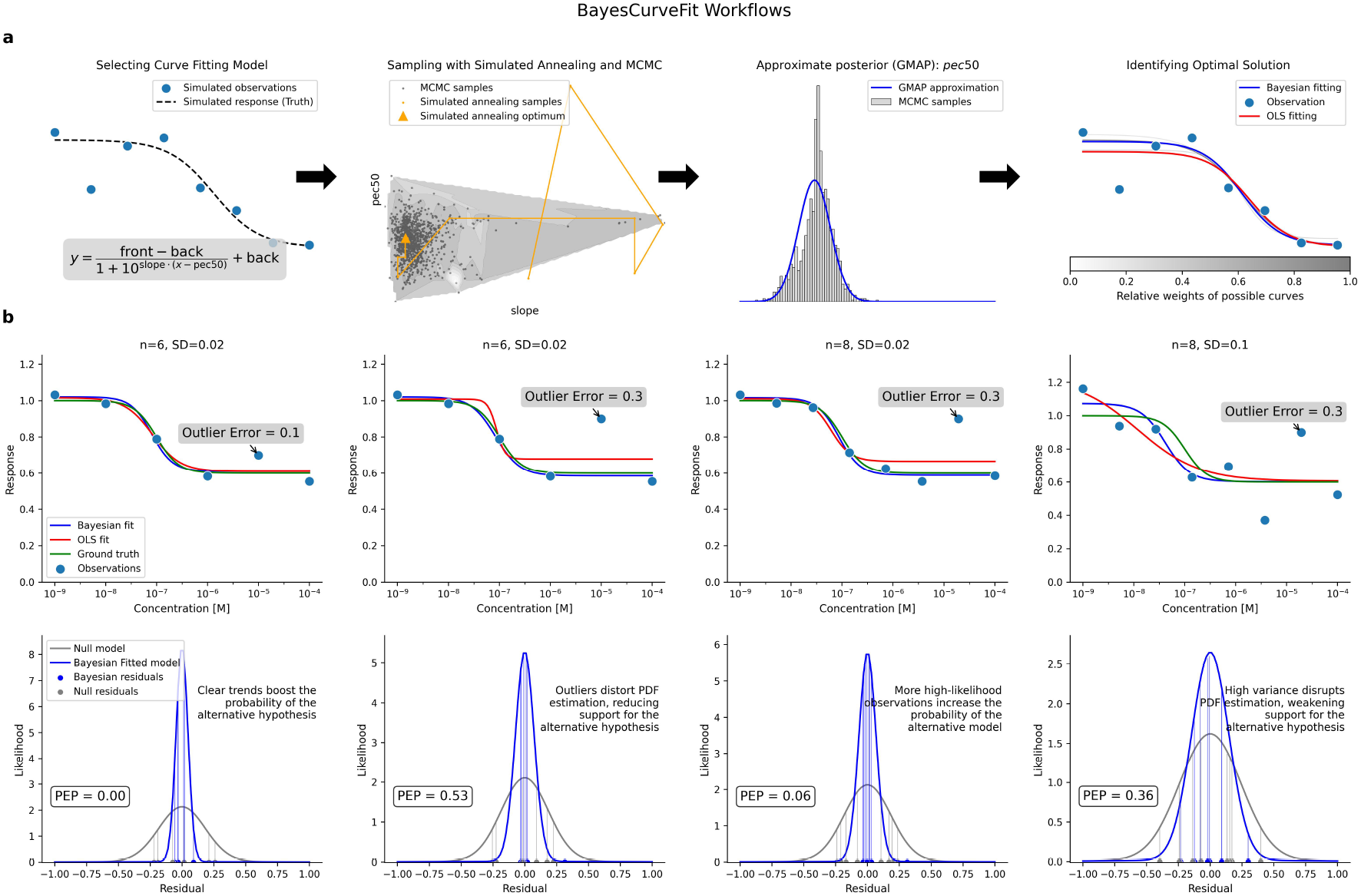
Overview of the *BayesCurveFit* workflow and its robustness to undersampled and noisy observations. (**a**) Schematic of the *BayesCurveFit* workflow. Users select a curve model (here, a four-parameter log–logistic function) to fit observed dose–response data. Global optimization via simulated annealing (orange trajectories) locates high-probability regions of the parameter space, followed by adaptive Markov chain Monte Carlo (MCMC) sampling to explore the posterior distribution under a Gaussian–Laplace mixture error model. The Gaussian Mixture Approximation to the Posterior (GMAP) integrates these samples, weighting each component by its posterior probability to produce correlation-aware parameter estimates and uncertainties. (**b**) Demonstration of *BayesCurveFit* performance under varying sample sizes, noise levels, and outlier magnitudes. Top panels compare *BayesCurveFit* (blue) and ordinary least squares (OLS; red) fits to the simulated ground truth (green). Gray lines indicate alternative posterior solutions weighted by their probabilities. Bottom panels show likelihood distributions of residuals under the null (gray) and alternative (blue) models; vertical lines denote observed residuals. The Posterior Error Probability (PEP) quantifies the probability that the observed data arise from noise. *BayesCurveFit* maintains accurate fits and stable PEP estimates even under outlier contamination and small-sample conditions.

### Step 1: Simulated Annealing for Initial Approximation

The inference begins with an initial posterior approximation obtained through Simulated Annealing (SA), a stochastic global optimization method that efficiently locates high-probability regions of parameter space. Simulated annealing balances exploration and exploitation by gradually lowering the search temperature, enabling the algorithm to escape local minima while converging toward stable solutions. This initialization provides an effective starting point for subsequent sampling, improving numerical stability and convergence efficiency. Prior Bayesian workflow analyses have shown that warm-started samplers typically yield faster mixing and better-calibrated posterior estimates than uninformed initializations [Yao et al., 2018].

### Step 2: Markov chain Monte Carlo for Posterior Sampling

From this initialization, we perform posterior sampling using an adaptive Markov chain Monte Carlo (MCMC) scheme formulated under a Gaussian–Laplace mixture error model. This hybrid error structure models the residuals as a weighted combination of Gaussian and Laplace components, providing flexibility to represent both moderate variance and infrequent large deviations in the data. The Gaussian component accommodates approximately homoscedastic noise, while the Laplace component models heavier-tailed behavior that may arise from outliers or heterogeneity among measurements. During sampling, adaptive updates of the proposal covariance and scaling factors maintain target acceptance rates and efficient exploration of the posterior space. The combined use of simulated annealing for initialization, adaptive MCMC for posterior exploration, and a Gaussian–Laplace mixture error model for likelihood evaluation defines the core sampling procedure applied across all experiments.

### Step 3: Gaussian Mixture Approximation for Estimating Uncertainty

To represent the posterior distribution compactly, we apply a Gaussian Mixture Approximation to the Posterior (GMAP). Rather than discrete Bayesian model averaging, GMAP fits a finite mixture of Gaussian components to the MCMC samples, providing a continuous, weighted approximation of the posterior density. This mixture representation preserves multimodal structure and variance heterogeneity while remaining analytically tractable for marginalization and prediction. Prior studies have demonstrated that Gaussian mixture approximations better capture uncertainty and prevent overconfident inference compared with single-mode estimates [Malsiner-Walli et al., 2017, Flock et al., 2025].

### Step 4: Measure Inference Reliability by Posterior Error Probability

For parameter-specific uncertainty assessment, we compute a Posterior Error Probability (PEP) for each fitted effect, defined as the probability that the observation belongs to the null or baseline distribution given the data and GMAP representation. PEP is computed from BIC (Bayesian Information Criterion) differences using the standard exp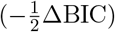 approximation to Bayes factors [Kass and Raftery, 1995], as detailed in **Supplementary Information (SI)**. PEP provides a probabilistic measure of inference reliability, analogous to the local false discovery rate but derived from the full posterior. This formulation aligns with prior large-scale Bayesian analyses where PEP offers interpretable and calibrated control of individual error rates [Käll et al., 2008, Shortreed et al., 2021].

These steps constitute a coherent Bayesian inference workflow. The design prioritizes stability, flexibility, and interpretability while maintaining compatibility with high-throughput data and heterogeneous noise typical of early screening experiments.

## 3 Results

The performance of *BayesCurveFit* is evaluated using both simulated and real datasets. Simulation studies are conducted to produce a representative of high-throughput screening conditions. We implemented a simulation module in BayesCurveFit to generate datasets based on user-specified fit functions, error probability density functions (PDFs), and parameter bounds.

In a real case study, we applied BayesCurveFit to identify drug-gene interactions in the classic Kinobeads dataset [Klaeger et al., 2017] and compared the results with those obtained from regression-based analyses in a state-of-the-art curve fitting method [Bayer et al., 2023].

### 3.1 Simulation benchmarking

For simulation benchmarking, we used the 40 *µ*M drug subset from the Cancer Therapeutic Response Portal (CTRP) [Seashore-Ludlow et al., 2015], containing 16 replicate dose–response measurements spanning concentrations from 1 × 10^−9^ M to 1 × 10^−4^ M. The previously published OLS (ordinary least squares)-fitted parameters from CurveCurator [Bayer et al., 2023] were treated as reference ground-truth values. Residuals from these OLS fits were extracted to estimate the empirical noise structure. The residual distribution of the CTRP dataset exhibited a long-tailed structure (**Supplementary Figure S1**). A two-component Gaussian mixture was therefore adopted to model both homoscedastic and long-tailed noise, serving as the empirical error distribution in subsequent simulations.

Synthetic dose–response curves were generated from the 4-parameter log–logistic function with 1,000 replicates per condition and sample sizes ranging from 6 to 50 measurement points. Random noise was added under two sampling schemes derived from the empirical residual model (**Supplementary Figure S1**): (i) **single-component sampling**, where errors were drawn solely from the dominant Gaussian component to approximate homoscedastic noise, and (ii) **full-mixture sampling**, where errors were drawn from the entire two-component Gaussian mixture to reproduce the empirically observed heavy-tailed variability.

Both *BayesCurveFit* and OLS were applied using identical parameter bounds to ensure a fair comparison. For each curve, the 4 log–logistic parameters (PEC_50_, slope, front, and back) were estimated, and accuracy was quantified using the normalized mean absolute error (MAE) across replicates—a scale-independent measure of deviation between estimated and true values.

Under the single-component noise condition (**Figure 2**, top row), all 3 methods showed similar convergence as sample size increased, with errors decreasing rapidly beyond 12 observations. Within the typical experimental range (6–12 points), *BayesCurveFit* achieved the highest accuracy for PEC_50_ and, more prominently, the slope parameter, whereas the front and back estimates were comparable across methods. The robust M-estimator did not confer a consistent advantage over OLS and, in several parameters, exhibited slightly higher error, suggesting that its loss reweighting provides limited benefit when noise is approximately Gaussian.

**Figure 2.**
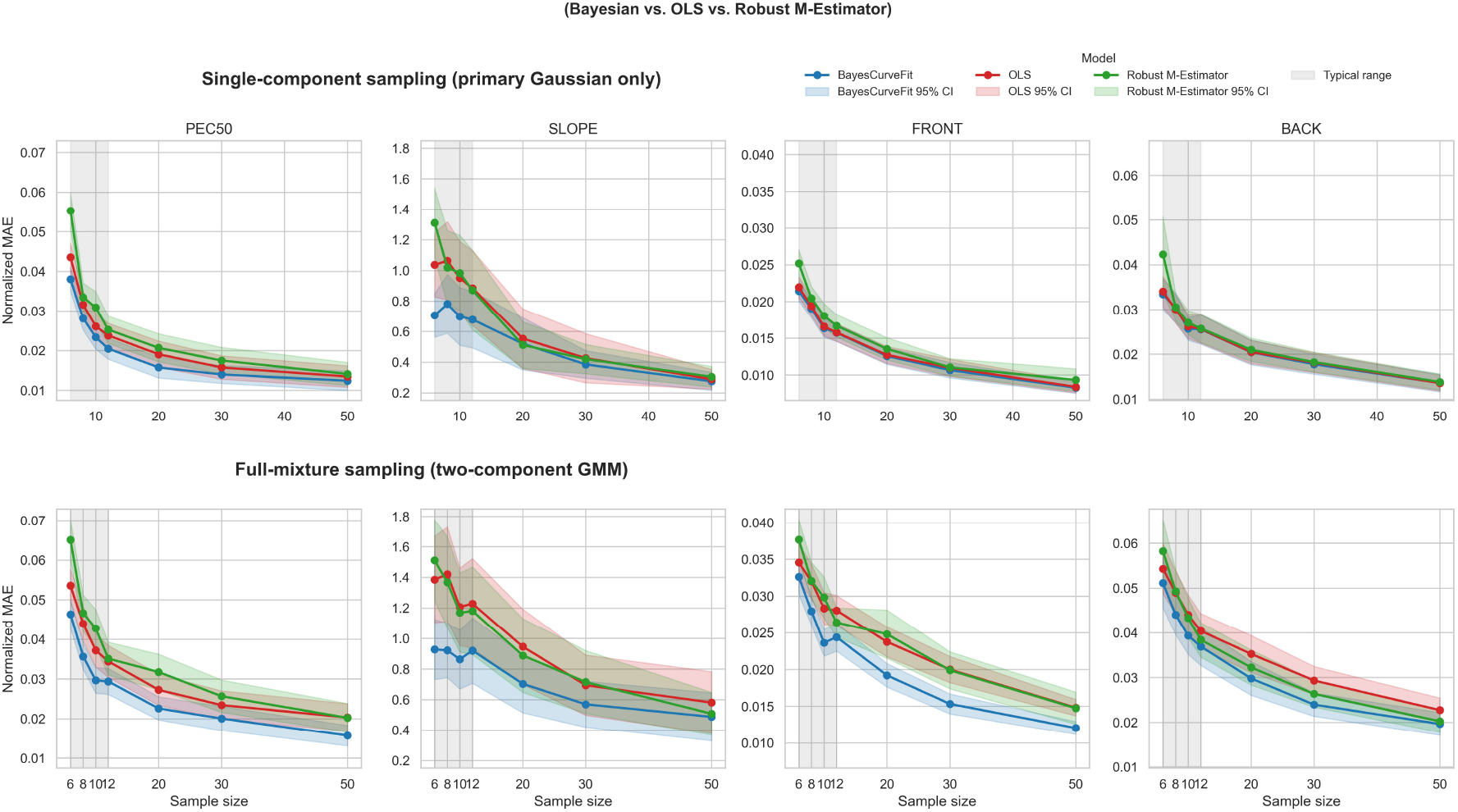
Simulation performance comparison of *BayesCurveFit*, OLS, and robust M-estimator fitting. Normalized mean absolute error (MAE) of fitted parameters (PEC_50_, slope, front, and back) as a function of sample size under two simulated noise regimes: *single-component sampling* (top; primary Gaussian only) and *full-mixture sampling* (bottom; two-component Gaussian mixture). Shaded regions indicate 95% confidence intervals across 1,000 simulation replicates. The gray band marks the typical experimental range (6–12 data points). *BayesCurveFit* (blue) consistently achieves the lowest estimation error and narrowest uncertainty bounds, particularly under heavy-tailed noise. The robust M-estimator (green) did not consistently outperform OLS (red) and in some parameters performed slightly worse, indicating that simple robust loss functions only partly mitigate heavy-tailed noise effects.

Under the full-mixture noise condition (**Figure 2**, bottom row), where heavy-tailed variability introduced occasional large deviations, all methods exhibited higher overall error. Nevertheless, *BayesCurveFit* remained the most resilient to this degradation, maintaining lower MAE values and narrower uncertainty bounds across parameters and sample sizes. The robust M-estimator sometimes performed worse than OLS in these settings, indicating that conventional robust regression can underperform when residuals are asymmetric or multimodal, conditions that are better handled by the Bayesian noise model.

Overall, these results demonstrate that *BayesCurveFit* provides more accurate and reliable parameter estimation than conventional OLS regression, particularly under small-sample and non-Gaussian noise conditions. Its robustness to experimental variability underscores its suitability for early-stage screening, where limited replication and uncontrolled noise often confound conventional curve-fitting approaches.

### 3.2 Experiments on real data

For the Kinobeads case study, dose–response measurements from 277 compounds across 54,223 protein targets were obtained from the CurveCurator data repository [Bayer et al., 2023]. The eight “Ratio” columns (Ratio 1–8) were used as dependent variables, and compound concentrations (1 × 10^−9^ M to 1 × 10^−4^ M) were defined as independent variables following the Kinobeads data format specification. Reference data for primary target validation were retrieved from the EMBL– EBI Open Targets platform [Ochoa et al., 2022] and the GtoPdb database [Harding et al., 2023].

We applied *BayesCurveFit* to the Kinobeads dataset previously analyzed by CurveCurator. This analysis compared PEP values generated by *BayesCurveFit* with the classical *p*-values reported by CurveCurator. To ensure methodological consistency, the log_2_ fold change (log_2_FC) was calculated using the same approach as in CurveCurator – based on model-predicted response ratios at the highest and lowest tested concentrations. This alignment allows a direct comparison between significance measures (PEP versus *p*-value) without introducing differences in fold-change computation.

Representative volcano plots for the drug *Imatinib* are shown in **Figure 3**. PDFGRB and KIT are the two well-known clinically validated primary targets [Apperley et al., 2002, Gajiwala et al., 2009, Cheah et al., 2014]. In the *BayesCurveFit* analysis, both candidates were clearly distinguished from background. However, neither the CurveCurator’s *p*-value-based analysis nor the original manual curation could classify either candidate as significant. This improvement in sensitivity primarily resulted from the use of PEP. Combined with log_2_FC estimates under long-tailed error distributions, PEP made these known target interactions readily discernible. Representative dose–response fits (**Figure 3 c–d**) further illustrate that *BayesCurveFit* preserved consistent monotonic response trends despite outlier measurements, whereas OLS-based fits in CurveCurator were distorted by the noise at extreme concentrations. Additional examples for *Lucitanib* and *LY-2584702* show similar improvements in signal–noise separation for their respective primary targets (**Supplementary Figure S2**), supporting the generality of *BayesCurveFit*.

**Figure 3.**
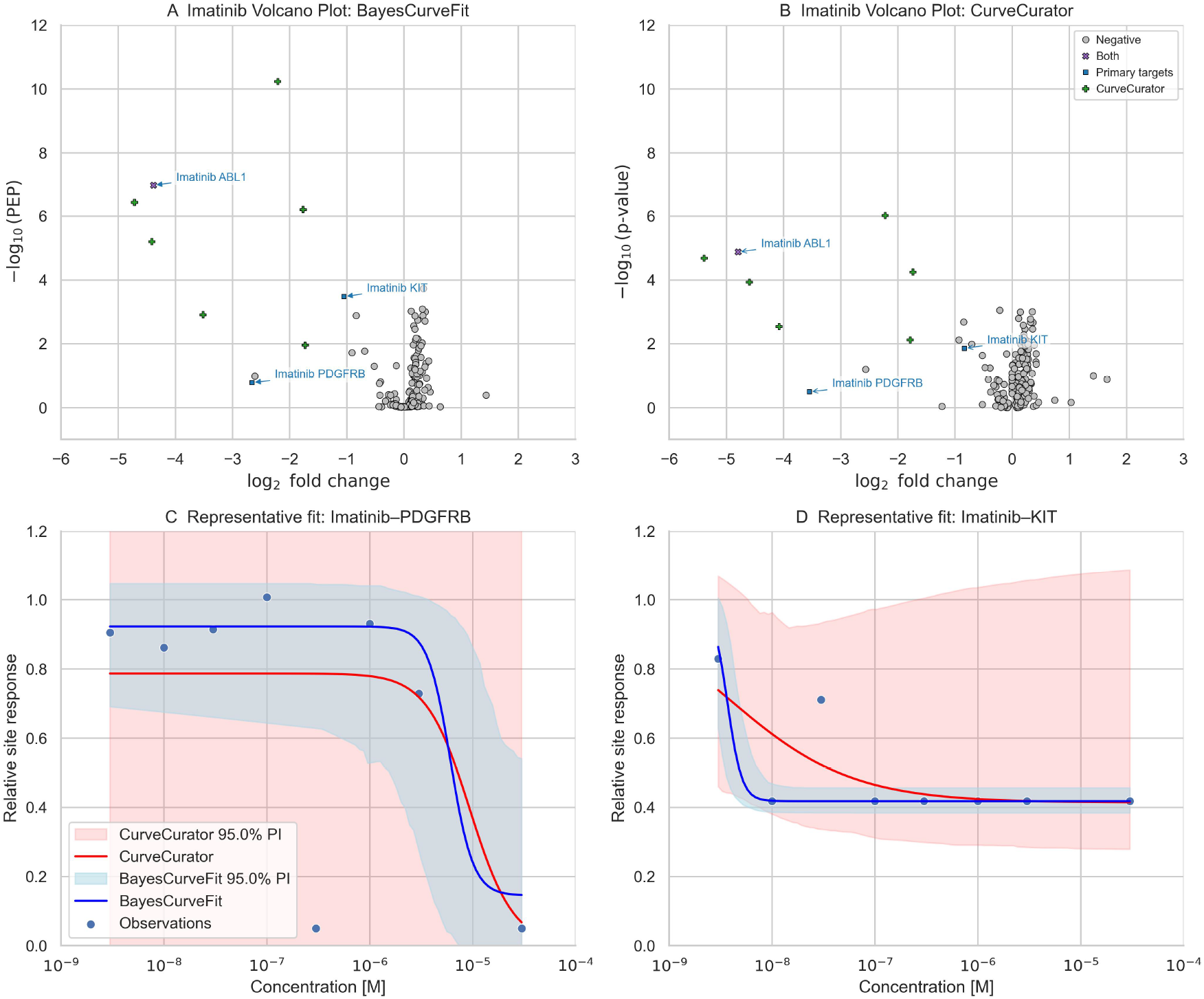
Comparison of Imatinib target detection using *BayesCurveFit* and CurveCurator. (a–b) Volcano plots for Imatinib showing log_2_ fold change versus significance metric (Bayesian posterior error probability, PEP, in *BayesCurveFit* ; and *p*-value in CurveCurator). PEP values were derived from Bayesian posterior inference in *BayesCurveFit*. PDGFRB and KIT, two well-established primary targets of Imatinib, were reported as non-significant by CurveCurator but showed markedly higher posterior probabilities of true response (lower PEP) in the *BayesCurveFit* analysis, indicating improved detectability under the Bayesian framework. Green plus markers denote targets identified as true regulations by CurveCurator, purple cross marks indicate primary targets identified by CurveCurator, and blue squares denote primary targets that were not identified by CurveCurator. (c–d) Representative dose–response fits for Imatinib–PDGFRB and Imatinib–KIT. Shaded regions indicate 95% credible (blue) or predictive (red) intervals. *BayesCurveFit* generates smoother, biologically plausible fits that remain stable despite outlier points, whereas OLS-based fits from CurveCurator are more influenced by noise at extreme concentrations.

To evaluate overall discriminative performance, we compared PEP- and *p*-value-based rankings across two reference sets: (i) the manual curation from the original Kinobeads study, which included all dose–response measurements, and (ii) known primary targets compiled from public databases, retaining only drugs with matching target annotations. The latter subset comprised 147 unique compounds and 29,317 compound–target records. To ensure numerical stability and a fair alignment with CurveCurator’s fitting procedure, parameter bounds in *BayesCurveFit* were defined using the same rule—anchored to each interaction’s observed response range—thereby minimizing boundary-related bias. Across both real benchmarks, *BayesCurveFit* consistently achieved higher areas under the ROC and precision–recall curves (**Figure 4**), indicating generally improved ranking of biologically validated interactions and reduced false-negative rates relative to *p*-value-based classification algorithms.

**Figure 4.**
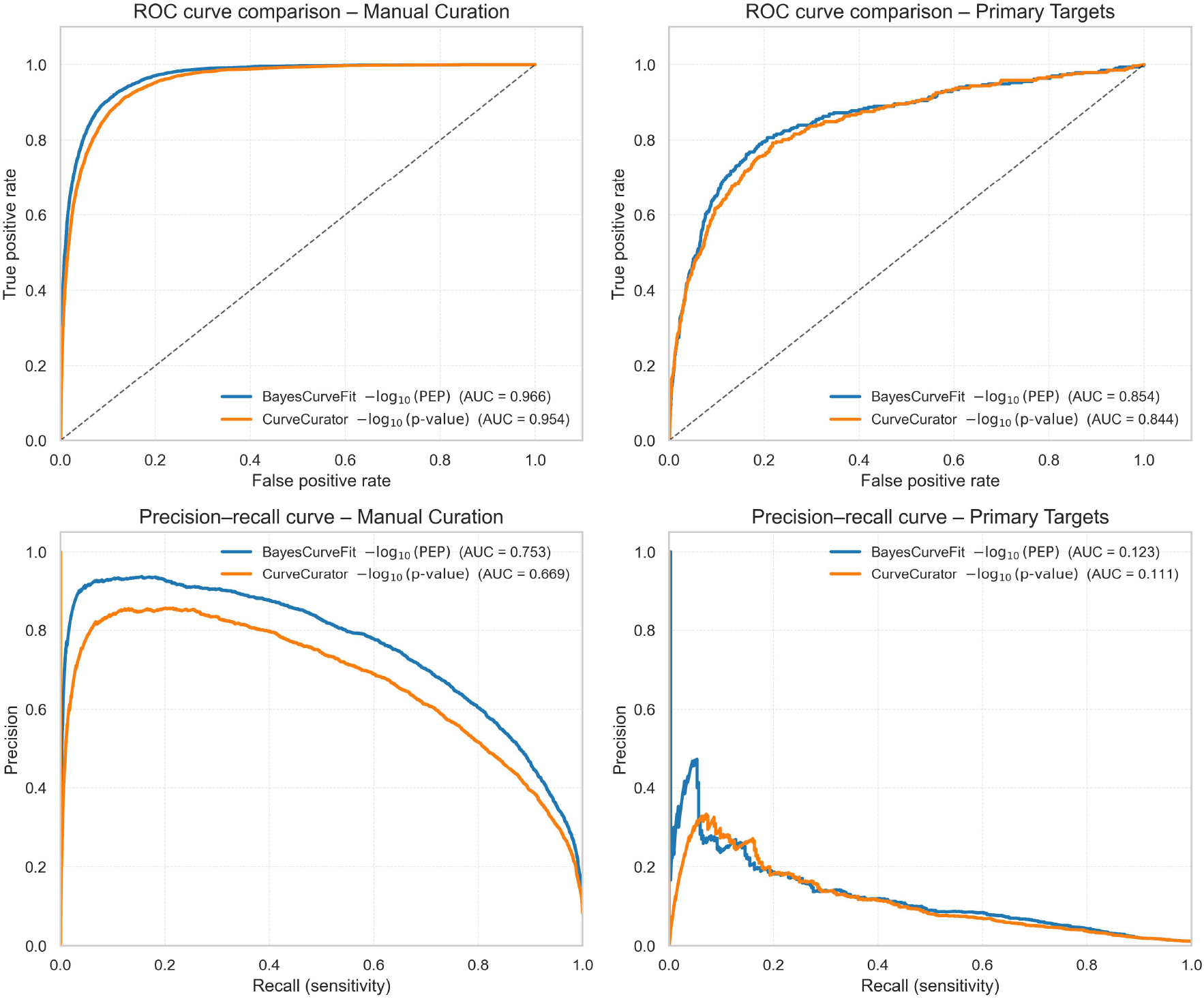
Performance comparison of *BayesCurveFit* and CurveCurator across all Kinobeads interactions. Receiver operating characteristic (ROC) and precision–recall (PR) curves comparing Bayesian PEP (blue) and classical *p*-value (orange) using manual curation (left) and primary target annotations (right) as ground truth. *BayesCurveFit* consistently achieves higher AUC values across both benchmarks, reflecting improved calibration and reduced false-negative rates for biologically validated targets.

## 4 Discussions

Accurate curve fitting under limited sampling and experimental noise remains a major challenge in early-stage drug discovery and other high-throughput assays [Inglese et al., 2007, Sebaugh, 2011]. A key advantage of *BayesCurveFit* lies in its robust parameter estimation compared with conventional regression-based approaches, particularly under data-limited conditions. This robustness is critical in practical applications where experimental throughput is limited by budget and the resulting error distributions often deviate from Gaussian assumptions. Our results show that the Gaussian–Laplace hybrid likelihood effectively accommodate long-tailed and asymmetric residuals, arising from plate effects and pipetting variability [Guan et al., 2023]. This resilience allows *BayesCurveFit* to recover true dose–response relationships that might otherwise be distorted or misclassified by traditional regression analyses.

Beyond parameter estimation, *BayesCurveFit* enhances the probabilistic interpretation of fit quality through PEP — a Bayesian analogue of the local false discovery rate [Käll et al., 2008]. Unlike a conventional *p*-value, PEP quantifies the probability that an observed response arises from noise, providing a direct measure of inference confidence. In the Kinobeads case study, the PEP-based framework is shown to enable clearer separation of true compound–target interactions from background variability, with the empirical comparison (**Supplementary Figure S3**) showing that PEP and nominal *p*-values diverge in sensitivity across the significance range. In general, PEP increases more gradually at low *p*, yielding conservative interpretation near traditional thresholds, but rises sharply in the high-*p* regime, improving discrimination among uncertain or inactive cases. Because the PEP provides a continuous and interpretable measure of confidence, it enables more rational adjustment of decision thresholds than fixed p-value cutoffs, mitigating the influence of minor fluctuations in noisy or data-limited settings.

Existing solutions based on robust loss functions partially mitigated the influence of outliers. However, they do not consistently outperform OLS, and often perform worse under asymmetric or multimodal noise—conditions frequently observed in high-throughput experiment data, which are better addressed by Bayesian frameworks that explicitly model uncertainty in the noise distribution [Gelman et al., 2013, Smid et al., 2020]. In simulation benchmarks, *BayesCurveFit* consistently achieved lower parameter estimation errors than both OLS and robust M-estimator baselines. Our simulations reproduced stochastic variability typical of high-throughput assays, including random, heteroscedastic, and heavy-tailed errors that often undermine deterministic regression under limited sampling. The two-component Gaussian mixture used for data generation was derived empirically from the existing CTRP residuals [Seashore-Ludlow et al., 2015] and applied identically across all methods, ensuring a fair comparison. During inference, *BayesCurveFit* used an independent Gaussian–Laplace noise model optimized within its own posterior framework, confirming that the observed performance gains arise from inferential robustness rather than model reuse or simulation bias. Structured artifacts such as hormesis, solubility limits, or compound degradation represent systematic deviations rather than stochastic variability. They are a separate set of problems and beyond the scope of this study, though they could potentially be addressed in extensions of *BayesCurveFit* through alternative curve formulations.

A Bayesian approach requires a specification of prior expectations, typically encoded through parameter bounds, which is a general limitation. If these bounds are too broad, convergence slows and posterior estimates become diffuse; when appropriately constrained, the sampler efficiently explores the parameter space and yields stable posteriors. In practice, we implemented data-dependent bounds for asymptotic parameters, constraining inference to empirically supported ranges and preventing extrapolation beyond observed responses (**Supplementary Information (SI)**: Dose–response equation and Initialization). Within these adaptive bounds, uniform priors act as weak regularization constraints rather than subjective assumptions, allowing the likelihood to dominate inference. Properly specified bounds promote efficient exploration and convergence, whereas overly diffuse limits tend to increase posterior variance and slow mixing, which can be assessed using standard convergence diagnostics (**Supplementary Information (SI)**: Gelman–Rubin 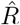, ESS).

*BayesCurveFit* is designed to address the challenges of data scarcity in early screening. It serves as a complement rather than a replacement for deterministic regression methods within drug discovery pipelines. Its probabilistic, inference-based framework is particularly valuable during exploratory phases, when replication is limited and validation resources are constrained. We recommend applying *BayesCurveFit* as a rescreening or prioritization tool for compounds initially classified as inactive. It can reveal important hits (e.g., *Imatinib*) that may have been overlooked due to data sparsity or technical noise. In this capacity, *BayesCurveFit* acts as a probabilistic safeguard, enhancing cost efficiency and decision reliability in the early stages of drug design.

## 5 Conclusion

Reliable inference of dose–response relationships from limited and noisy experimental data remains a persistent challenge in early-stage drug discovery. To address this gap, we developed *BayesCurveFit*, a Bayesian framework for dose–response curve fitting that integrates stochastic optimization and MCMC-based posterior sampling within a unified inference pipeline for compound screening. Using both simulated datasets and large experimental benchmarks, we demonstrate that *BayesCurveFit* substantially improves parameter recovery and target ranking accuracy compared with conventional ordinary least squares (OLS) regression. Furthermore, the inclusion of posterior error probability (PEP) provides a coherent probabilistic measure of confidence, enabling interpretable discrimination between true and spurious dose–response patterns.

The principles underlying *BayesCurveFit* are broadly applicable. Its probabilistic inference, adaptive noise modeling, and modular Bayesian architecture naturally extend to other quantitative biology contexts, including dose–toxicity assessment, omics-based benchmark dose estimation, and other analyses where uncertainty is intrinsic and data are limited. We have implemented *BayesCurveFit* as an open-source Python package to make advanced Bayesian modeling readily accessible within practical bioinformatics workflows.

## 6 Code and Data Availability, and Acknowledgment

The BayesCurveFit software (both source code and a Python package that can be automatically installed via pip) and documentations are freely available on GitHub under the open source Apache 2.0 license: https://github.com/ndu-bioinfo/BayesCurveFit. The version of BayesCurveFit used in this manuscript is v0.6.5.

Computational experiments and analyses were conducted on a workstation running Ubuntu 22.04.5 LTS (Jammy Jellyfish) with the Linux kernel 6.8.0-78-generic on an x86 64 architecture. Installation instructions, example usage, and additional experiment results can be found in **Supplementary Information (SI)**. Datasets and Jupyter Notebooks for reproducing experiment results in Section 3 can be found at: https://github.com/ndu-ioinfo/bayescurvefit notebooks

This work is partially supported by the National Institute of Allergy and Infectious Diseases (NI-AID) of the US National Institutes of Health (NIH) under grant numbers P01AI168347, R01AI184931, and U01AI187057.

