## Supplementary Document for "BayesCurveFit: Enhancing Curve Fitting for Early-Stage Compound Screening Using Bayesian Inference"

#### Supplementary Information Overview

This document provides supplementary figures, detailed implementation notes, and methodological derivations supporting the main text. Section A presents additional figures referenced in main text. Section B provides installation and usage instructions with a reproducible example. Section C details the full inference pipeline, including the mathematical formulations, Bayesian model averaging, posterior error probability derivation, and convergence diagnostics.

Code and Documentations of BayesCurveFit:

https://github.com/ndu-bioinfo/BayesCurveFit

Data and Jupyter Notebooks for reproducing analysis results including Figures in this manuscript:

https://github.com/ndu-ioinfo/bayescurvefit_notebooks

### A Supplementary Figures

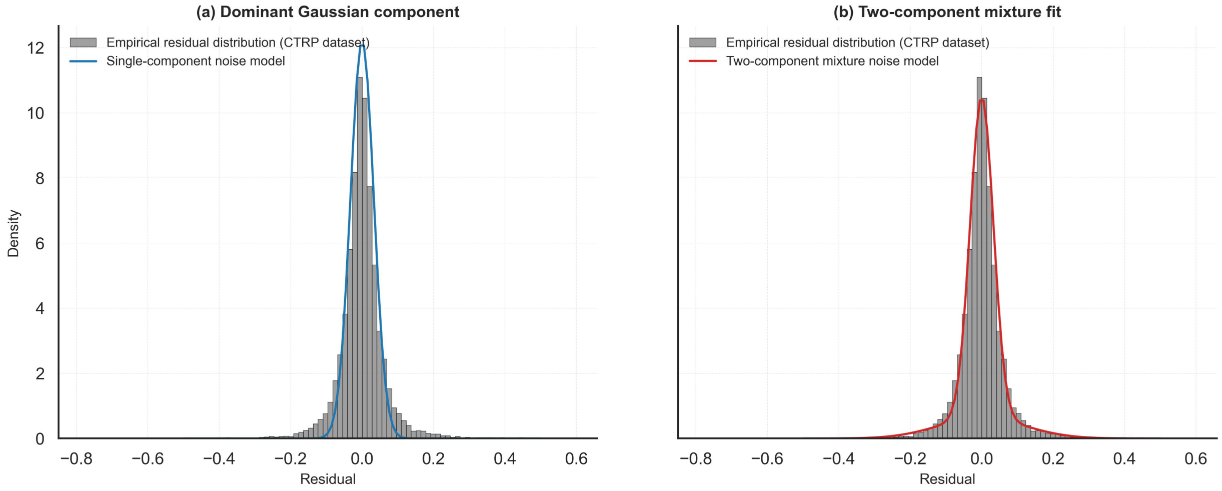

Figure S1: **Residual distribution of CTRP data and noise models used for simulation.** Gray bars represent the empirical residuals extracted from OLS fits of the CTRP dataset. (a) Single-component noise model derived from the dominant Gaussian component, representing homoscedastic experimental errors. (b) Two-component mixture noise model capturing both central and long-tailed variability observed in the residuals. The mixture model was used as the empirical error distribution in subsequent simulation studies.

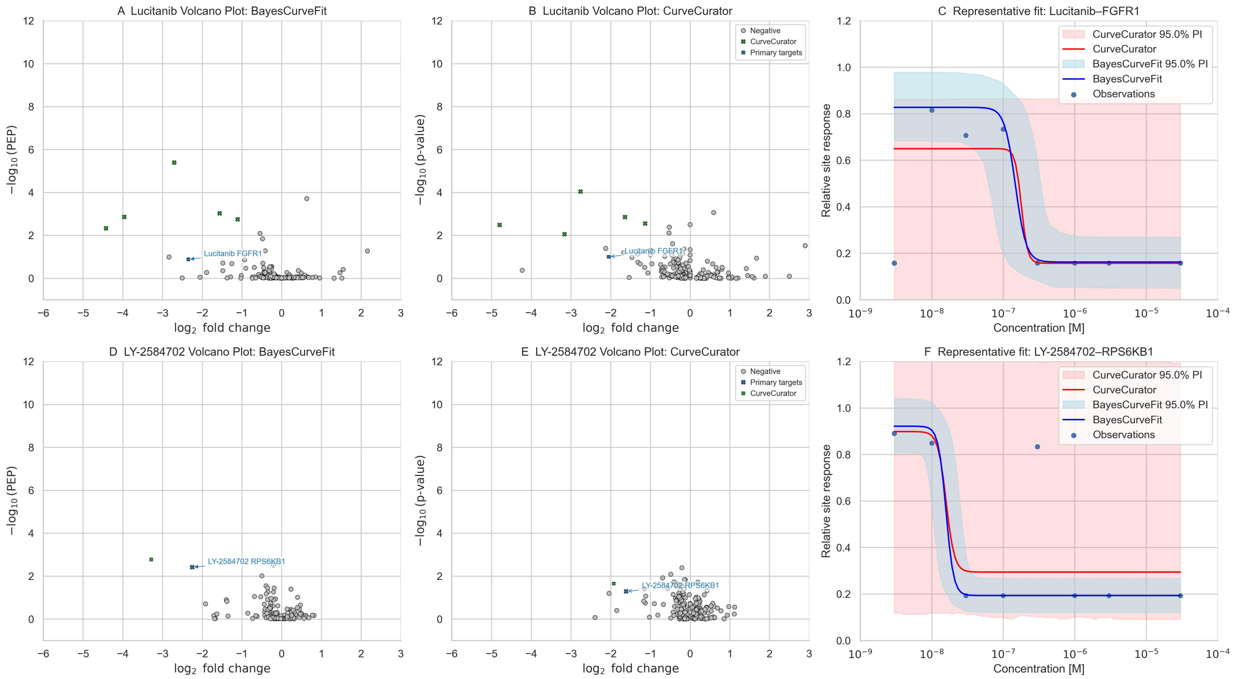

Figure S2: **Additional examples demonstrating improved signal–noise separation using *BayesCurveFit*.** (a–c) Volcano plots and representative dose–response fits for Lucitanib. The primary target FGFR1 displays clearer separation from background interactions in the *BayesCurveFit* analysis compared to CurveCurator. (d–f) Volcano plots and representative fits for LY-2584702. The downstream target RPS6KB1 similarly shows improved distinction from noise, with *BayesCurveFit* maintaining a stable, monotonic response curve and tighter uncertainty bounds. In both examples, the Bayesian posterior provides improved discrimination of biologically meaningful responses from background variability, consistent with the results observed for Imatinib. Green plus markers denote targets identified as true regulations by CurveCurator, purple crosses indicate primary targets identified by CurveCurator, and blue squares denote primary targets not identified by CurveCurator. Shaded regions represent 95% credible (blue) and predictive (red) intervals for *BayesCurveFit* and CurveCurator fits, respectively.

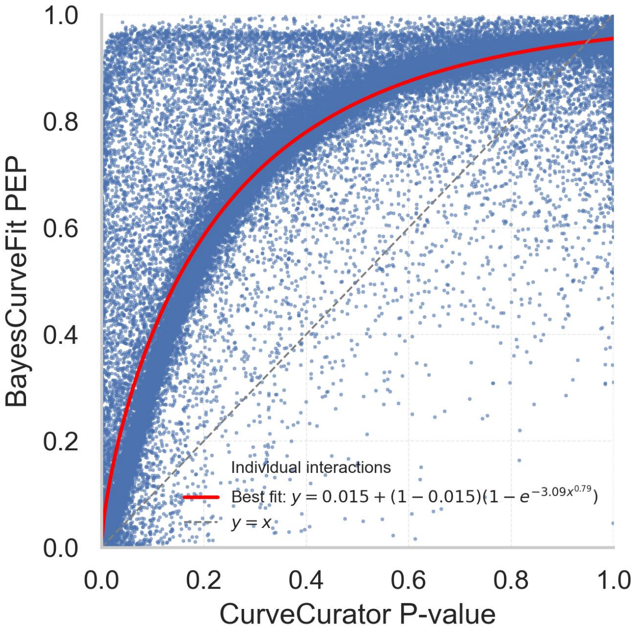

Figure S3: **Relationship between Bayesian posterior error probability (PEP) and nominal *p*-values.** Each point represents a single compound–target fit. The solid red line shows the best-fit exponential saturation model ($y = 0.015 + (1 - 0.015)\left[1 - e^{-3.09x^{0.79}}\right]$), capturing the nonlinear correspondence between Bayesian and frequentist significance measures. The dashed gray line denotes the identity line ($y = x$). A nominal $p$-value of 0.05 corresponds approximately to a PEP of 0.2, illustrating that Bayesian posterior probabilities yield more conservative significance estimates relative to classical $p$-values.

### B Installation and Example Usage

#### B.1 Installation

*BayesCurveFit* is a modular Python library designed for both standalone use and integration into existing workflows. It can be installed via:

```
pip install bayescurvefit
```

or by running the `make` command as provided in the source repository: https://github.com/ndu-bioinfo/BayesCurveFit

#### B.2 Running an Example

**Note:** Due to platform dependency of the underlying MCMC process, small numerical differences may occur across operating systems or architectures. These variations are inherent to stochastic sampling and do not affect result validity.

Input

```python
import numpy as np
from bayescurvefit.execution import BayesFitModel

# Define the four-parameter log-logistic function
def log_logistic_4p(x, pec50, slope, front, back):
    return (front - back) / (1 + 10 ** (slope * (x + pec50))) + back

x_data = np.array([-9.0, -8.3, -7.6, -6.9, -6.1, -5.4, -4.7, -4.0])
y_data = np.array([1.12, 0.74, 1.03, 1.08, 0.76, 0.61, 0.39, 0.38])

params_range = [(5, 8), (0.01, 10), (0.28, 1.22), (0.28, 1.22)]
param_names = ["pec50", "slope", "front", "back"]

# Run the Bayesian fitting pipeline
run = BayesFitModel(
    x_data=x_data,
    y_data=y_data,
    fit_function=log_logistic_4p,
    params_range=params_range,
    param_names=param_names,
)
run.get_result()
```

#### B.3 Output Interpretation

Parameter names prefixed with `fit_` and `std_` represent posterior means and standard deviations, respectively. The posterior error probability (`pep`) quantifies the Bayesian probability that the observed effect arises from noise. A low PEP indicates strong evidence for a true dose–response relationship.

Output

```
fit_pec50   5.856   std_pec50   0.185
fit_slope   1.089   std_slope   0.399
fit_front   1.063   std_front   0.062
fit_back    0.377   std_back    0.051
est_std     0.090   rmse        0.123
pep         0.063   convergence_warning False
```

### C Method Details for Pipeline Implementation

#### C.1 Dose Response Equation

The four-parameter log-logistic equation (1) is a commonly used model for describing dose-response relationships in pharmacology and toxicology. It characterizes how a biological response changes with varying concentrations of a substance, such as a drug or a toxin. The equation is expressed as:

$$y = \frac{front - back}{1 + 10^{slope \cdot (x - pec50)}} + back \tag{1}$$

where:

- $y$ is the ratio of response to the reference.
- $front$ is the upper asymptote.
- $back$ is the lower asymptote.
- $slope$ is the slope factor.
- $pec50$ is the $-\log_{10}$ transformed dose at which the response is halfway between $front$ and $back$.
- $x$ is the $-\log_{10}$ transformed dose concentration.

#### C.2 Null Model

The null model serves as a baseline for comparison with more complex models. It assumes that the dependent variable is independent of any predictors. Mathematically, the null model (4) is:

$$f(x) = c + \epsilon \tag{2}$$

where $f(x)$ is the predicted value, a constant $c$, and $\epsilon$ represents the error term or random noise centered at zero, which accounts for any variability not explained by the constant. This model suggests that the outcome does not vary with the input $x$ and remains constant on average, with deviations due to random errors.

#### C.3 Initial Approximation and Posterior Sampling

Upon loading the data, *BayesCurveFit* first employs Simulated Annealing (SA) to estimate the point-wise global optimum $\hat{\theta}$ and the nuisance parameters of the probability density function (PDF). In its default configuration, the algorithm jointly estimates the parameters $\theta$ and the standard deviation $\sigma$ of the Gaussian component by maximizing the conditional posterior probability:

$$(\hat{\theta}, \hat{\sigma}) = \arg\max_{\theta,\sigma} \left\{ P(\theta, \sigma \mid y) \propto P(y \mid \theta, \sigma)\, P(\theta)\, P(\sigma) \right\}. \tag{3}$$

Here, $\hat{\theta}$ and $\hat{\sigma}$ denote the maximum a posteriori (MAP) estimates of the parameters. The posterior probability $P(\theta, \sigma \mid y)$ is proportional to the product of the likelihood $P(y \mid \theta, \sigma)$ and the priors $P(\theta)$ and $P(\sigma)$. While no specific prior distributions are imposed, users can define practical parameter bounds that act as uniform priors. For $\theta$, these bounds restrict sampling to physically or biologically reasonable regions. For $\sigma$, the range is automatically set between $1 \times 10^{-6}$ and the standard deviation of all observations, ensuring that the fitted variance remains below the total observed noise. Within these boundaries, the likelihood function primarily drives the optimization.

The initial optimization employs a Gaussian likelihood solely as a numerically stable approximation for the error term. This choice ensures smooth exploration of the parameter space during simulated annealing (SA), where discontinuities in non-Gaussian likelihoods can otherwise hinder convergence. Importantly, this Gaussian assumption is used only for initialization: subsequent posterior inference adopts a Gaussian–Laplace hybrid error model that captures non-Gaussian features such as heavy tails and outlier-prone residuals, allowing realistic modeling of non-Gaussian noise.

A null model, assuming $y$ is independent of $x$, is also fitted using SA with a fixed $\sigma$ equal to the standard deviation of all observations. This null model provides a baseline for later false positive estimation through posterior error probability (PEP) computation.

Following the SA-based initialization, *BayesCurveFit* performs Markov Chain Monte Carlo (MCMC) sampling to characterize the posterior distribution of the parameters. Each chain is initialized from the global optimum determined by SA and extended by 1,000 steps per iteration. Sampling continues until the Gelman–Rubin diagnostic and Effective Sample Size (ESS) criteria [Gelman et al., 2013] indicate convergence or until the maximum iteration limit is reached. If convergence is not achieved, a warning is flagged in the final output for that curve.

MCMC execution is implemented using the lightweight `emcee` package [Foreman-Mackey et al., 2019], which offers efficient parallel sampling. By default, *BayesCurveFit* uses the *DESnookerMove* algorithm for parameter spaces with more than three dimensions [Braak and Vrugt, 2008] and *StretchMove* for lower-dimensional problems [Goodman and Weare, 2010]. To prevent premature divergence from the global optimum, a weak Gaussian penalty term is incorporated into the log-likelihood function:

$$\text{penalties} = -0.5 \left( \frac{\|\theta_{\text{proposed}} - \hat{\theta}\|_2}{\sigma_{\text{penalty}}} \right)^2 . \tag{4}$$

This penalty is mathematically equivalent to imposing a shallow Gaussian prior centered on the simulated annealing optimum $\hat{\theta}$. It acts as a local stabilizer that constrains early MCMC proposals to plausible regions of parameter space while still allowing full exploration as the chain progresses. The scaling factor $\sigma_{\text{penalty}} = 3.0$ corresponds to approximately a 99.7% confidence interval under a standard normal distribution, ensuring that the sampler remains near the global optimum unless substantial evidence supports an alternative high-probability region.

#### C.4 Gaussian Mixture Approximation to the Posterior (GMAP)

After the posterior samples are obtained via MCMC, the marginal posterior distribution is approximated using a Gaussian Mixture Model (GMM). To smooth sampling noise and preserve multimodality, a kernel density estimate (KDE) is first applied to resample the posterior space, after which candidate GMMs with one to ten components ($K = 1, \ldots, 10$) are fitted. The optimal number of mixture components is selected automatically using an information-criterion-based approach (Bayesian Information Criterion, introduced in the next step), which balances model fit and complexity to prevent overfitting. Note, here the BIC refers to the internal criterion computed by the GMM during fitting to select the most parsimonious posterior representation; this is distinct from the downstream BIC calculation later used for model comparison. The selected GMM thus provides a data-driven and parsimonious approximation of the full posterior distribution, preserving both parameter covariance and potential multimodal structure.

The joint probability density of all parameters $\theta$, corresponding to each Gaussian component in the multivariate GMM, is expressed as:

$$P(\theta \mid \mathbf{y}) \approx \sum_{k=1}^{K} \pi_k \, \mathcal{N}(\theta \mid \mu_k, \Sigma_k), \tag{5}$$

where $K$ is the number of mixture components, $\pi_k$ are the mixture weights, $\mu_k$ are the component mean vectors, and $\Sigma_k$ are the corresponding covariance matrices.

The mixture-weighted posterior mean and covariance summarize the parameter distribution implied by the fitted GMM:

$$\bar{\theta} = \sum_{k=1}^{K} \pi_k \, \mu_k, \tag{6}$$

and the corresponding covariance matrix, capturing both within-component uncertainty and between-component variability, follows from the law of total variance:

$$\Sigma_\theta = \sum_{k=1}^{K} \pi_k \left( \Sigma_k + (\mu_k - \bar{\theta})(\mu_k - \bar{\theta})^\top \right). \tag{7}$$

The posterior standard deviation of parameter $\theta_i$ given the data $\mathbf{y}$ is then:

$$\sigma(\theta_i \mid \mathbf{y}) = \sqrt{\bar{\Sigma}_{ii}}. \tag{8}$$

The overall posterior density is approximated by a Gaussian mixture, combining component-wise contributions weighted by their respective mixture responsibilities:

$$\log p(\theta \mid \mathbf{y}) \approx \log \sum_{k=1}^{K} \pi_k \, \mathcal{N}(\theta;\, \mu_k,\, \Sigma_k)\,. \tag{9}$$

This formulation represents the posterior probability density under the Gaussian Mixture Approximation to the Posterior (GMAP), combining contributions from all local modes weighted by their posterior probabilities. Each component $\mathcal{N}(\theta; \mu_k, \Sigma_k)$ captures both the location and covariance structure of a local mode, and the mixture weights $\pi_k$ reflect the relative evidence for each mode.

From this fitted mixture, we approximate the model evidence by evaluating the overall likelihood of the observed data under the posterior mixture. Rather than deriving this quantity analytically from

Bayes' rule, we compute an empirical, posterior-weighted estimate of the likelihood by evaluating the data likelihood at the mean of each mixture component and aggregating the results according to the fitted mixture weights:

$$\log \tilde{L}(\mathbf{y}) = \log \sum_{k=1}^{K} \pi_k \, p(\mathbf{y} \mid \mu_k) \,, \tag{10}$$

where $p(\mathbf{y} \mid \mu_k)$ denotes the likelihood of the observed data evaluated at the mean of each mixture component. This mixture-integrated likelihood provides a practical and numerically stable approximation to the marginal model evidence by averaging over all posterior modes, and is evaluated using the Gaussian mixture model with the optimal number of components $K$ selected by the internal BIC criterion during fitting.

#### C.5 Calculating Posterior Error Probability (PEP)

Model evidence is then quantified using the Bayesian Information Criterion (BIC) [Kass and Raftery, 1995]:

$$\text{BIC} = p \log N - 2 \log \tilde{L}(\mathbf{y}), \tag{11}$$

where $p$ is the number of fitted parameters and $N$ is the number of observations. In the classical formulation, the likelihood term $p(\mathbf{y} \mid \hat{\theta})$ is evaluated at the maximum-likelihood estimate $\hat{\theta}$. In our GMAP implementation, this term is replaced by the posterior-weighted likelihood $\tilde{L}(\mathbf{y})$, which integrates contributions from all mixture components of the posterior approximation (Eq. 9).

This mixture-integrated form smooths numerical fluctuations arising from multimodal or noisy posteriors and complements the penalized Gaussian sampling strategy used during posterior estimation—both approaches aim to stabilize inference by preventing overfitting to narrow local modes and by preserving multiple plausible solutions when supported by the data. This substitution preserves the BIC's model complexity penalty while providing a more stable, mixture-based estimate of model evidence.

Together, these steps provide a numerically stable approximation to the marginal model evidence, although this formulation is not strictly identical to the classical BIC definition based on a single maximum-likelihood estimate.

For model selection (e.g., determining the optimal number of mixture components), the dominant mixture component ($\mu_{k^*}$, corresponding to the highest-weight or maximum-likelihood mode) can be used as a plug-in estimate of $\bar{\theta}$, yielding rankings that are expected to be consistent with the mixture-integrated form when the posterior approximation is unimodal or highly concentrated.

To compare the alternative model against the null hypothesis, we first compute the relative information loss for each model based on its BIC value:

$$\Delta_0 = \text{BIC}_0 - \min(\text{BIC}_0, \text{BIC}_1), \tag{12}$$

$$\Delta_1 = \text{BIC}_1 - \min(\text{BIC}_0, \text{BIC}_1), \tag{13}$$

where $\text{BIC}_0$ and $\text{BIC}_1$ denote the null (no relationship between $x$ and $y$) and alternative (fitted) models, respectively. Smaller $\Delta$ values indicate stronger relative support for a given model, with $\Delta = 0$ identifying the best-fitting one.

The posterior error probability (PEP) is then derived from these BIC differences as:

$$PEP = \frac{\exp(-0.5\Delta_0)}{\exp(-0.5\Delta_0) + \exp(-0.5\Delta_1)}. \tag{14}$$

The expression in Eq. (14) follows directly from the relationship between BIC differences and Bayes factors [Kass and Raftery, 1995]. Specifically, $\exp(-0.5\Delta)$ approximates the marginal likelihood ratio between competing models. Normalizing these quantities across the null and alternative models yields the posterior probability of the null model given the data—defined here as the PEP. This formulation therefore converts BIC-derived model evidence into an interpretable probabilistic measure of model plausibility.

Conceptually, the PEP represents the probability that the observed data are better explained by random variation than by a true dose–response relationship. In practice, it serves as a Bayesian analogue to the local false discovery rate, providing a continuous measure of evidence against the null model rather than a binary hypothesis decision. Because it is derived directly from model probabilities, PEP avoids reliance on asymptotic $p$-value assumptions and remains interpretable even under non-Gaussian or small-sample conditions.

#### C.6 Nuisance Parameter Fine Tuning

Accurately estimating the true $\sigma$ in these scenarios is challenging because we are uncertain if the under-weighted observations are outliers or valid observations. However, the real $\sigma$ value must lie between the estimated $\sigma$ and the RMSE. A simplified solution is to use the mean of these two values as the estimate of the true $\sigma$. In cases where $\sigma$ is fitted to errors without biased fitting, the $\sigma$ estimate will be almost equal to the calculated RMSE. Therefore, in either case, $\sigma$ can be estimated by taking the mean of the initial $\sigma$ estimation and the corresponding RMSE. The function (5) for the adjusted $\sigma$ is given by:

$$\sigma_{\text{adj}} = \frac{\sigma_{\text{approx}} + \text{RMSE}}{2} \tag{15}$$

where $\sigma_{\text{approx}}$ is the initially approximated $\sigma$, and RMSE is the Root Mean Square Error of the observation residues. Finally, we re-run the SA step to improve the $\theta$ estimation using the MAP approach with the PDF incorporating the adjusted $\sigma$.

#### C.7 Gelman-Rubin Convergence Diagnostic

The Gelman-Rubin diagnostic, also known as the R-hat statistic, is employed to assess the convergence of MCMC simulations. This method evaluates whether multiple chains have converged to a common stationary distribution, indicating that the chains have mixed well and are providing reliable samples from the posterior distribution. To enhance the accuracy of the R-hat statistic, each MCMC chain is split into two halves [Gelman et al., 2013] to ensure the within-chain mixing is better represented at different times.

Following this, the between-chain variance $B$ (6) and within-chain variance $W$ (7) are calculated as follows:

$$B = \frac{n}{m-1} \sum_{j=1}^{m} (\bar{\theta}_{\cdot j} - \bar{\theta}_{\cdot\cdot})^2 \tag{16}$$

$$W = \frac{1}{m} \sum_{j=1}^{m} s_j^2 \tag{17}$$

where $n$ is the number of samples per chain, $m$ is the number of chains, $\bar{\theta}_{\cdot j}$ is the mean of the $j$-th chain, $\bar{\theta}_{\cdot\cdot}$ is the mean of the means of all chains, and $s_j^2$ is the variance within the $j$-th chain.

The total variance $\widehat{\text{Var}}^+$ (8) is estimated as:

$$\widehat{\text{Var}}^+ = \left(1 - \frac{1}{n}\right) W + \frac{1}{n} B \tag{18}$$

And the R-hat statistic (9) is then calculated as:

$$\hat{R} = \sqrt{\frac{\widehat{\text{Var}}^+}{W}} \tag{19}$$

The R-hat provides an estimate of the potential scale reduction factor, which decreases to 1 as $n \to \infty$. Therefore, the closeness of the R-hat values to 1 indicates that the chains have probably converged, and we selected 1.1 as the maximal threshold for R-hat to claim convergence.

#### C.8 Effective Sample Size (ESS)

The ESS [Gelman et al., 2013] measures the number of independent samples equivalent to the correlated samples produced by MCMC simulations. The ESS helps determine the quality of the samples and the reliability of the parameter estimates. To calculate ESS, we followed the methodology described by Gelman et al. [2013]. The process begins with computing the variogram $V_t$ (10) at each lag $t$ to understand the autocorrelation within the MCMC chains:

$$V_t = \frac{1}{m(n-t)} \sum_{j=1}^{m} \sum_{i=1}^{n-t} (\theta_{i+t,j} - \theta_{i,j})^2 \tag{20}$$

where $\theta_{i,j}$ represents the $i$-th sample in the $j$-th chain, $m$ is the number of chains, and $n$ is the number of samples per chain. The ESS (11) is then calculated based on the variogram and total variance computed in Equation 18:

$$\text{ESS} = \frac{n \cdot m}{1 + 2\sum_{t=1}^{T} \hat{p}_t} \tag{21}$$

The parameter $T$ is chosen as the first lag where the mean autocorrelation across all parameters falls below a threshold, where $n$ is the number of samples per chain, $m$ is the number of chains, and $\hat{p}_t$ (12) is the estimated autocorrelation at lag $t$:

$$\hat{p}_t = 1 - \frac{V_t}{2\widehat{\text{Var}}^+}. \tag{22}$$

We set the minimal threshold for ESS to 100 for each parameter. According to Gelman et al. [2013], 100 independent draws ensure that the MCMC error adds negligible uncertainty to the actual posterior variance.

#### C.9 Non-GMAP Adjustment

The GMAP approach, while powerful, is slow because it requires MCMC posterior sampling. Some users may consider it is not always worthwhile to run a GMAP process in scenarios when the parmeter uncertainty is low and only a rough estimate of the fitting is needed. Moreover, GMAP assumes that the uncertainties follow a normal distribution, which might require transformations of parameters before running the BayesCurveFit pipeline. For instance, the ec50 value was -log10 transformed to pec50 in Equation 1. Some users do not prefer to transform the data. For all these users, we offer an alternative OLS-based correction option without using MCMC. The bottom line is the BayesCurveFit-generated fitting results are at least as good as the best the OLS fitting could achieve.

The optinal Non-GMAP workflow in BayesCurveFit consists of two steps. Firstly, an OLS fitting is attempted using the `curve_fit` function from the `scipy.optimize` module, with bounds identical to the user-defined bounds and without an initial estimate. If the regression fails to converge, a second attempt is made using the SA-determined optima as initial estimations. The OLS fitted result is then compared to the SA optima based on their log likelihoods, and the final optima are determined by whichever has the higher log likelihood.

The simulated annealing (SA) approach in BayesCurveFit is used to approximate the global optima. It may terminate early when the peak of the parameter's posterior distribution is not sharp enough, where the difference between two sampled points is less than the required minimal log probability difference. This issue could potentially be solved by setting an extremely low delta threshold, at the cost of increased computation time, particularly when the temperature decreases in the later stages of the SA process. Therefore, we implemented the OLS adjustment option to effectively help determine the optima, leading to an optimal detection that is not worse than OLS by itself, particularly in cases when the OLS process is hard to converge.

One limitation of the point-wise approximation is that the likelihood value may be inflated because it does not account for suboptimal solutions, as discussed in the MCMC sampling method. To address parameter uncertainties without fully sampling the posterior space, we employed an approximation approach modified from leave-one-out (LOO) cross-validation [Gelman et al., 2013].

Specifically, we use the approximated global optima, denoted as $\hat{\theta}$, as initial parameters and rerun the OLS fitting for each 'training' LOO $(x^{(i)}_{\text{train}}, y^{(i)}_{\text{train}})$ pair. In each iteration $i$, one observation $(x_i, y_i)$ is temporarily excluded from the training set to form the LOO training pair $(x^{(i)}_{\text{train}}, y^{(i)}_{\text{train}}) = \{(x_j, y_j) \mid j \neq i\}$. This resampling procedure allows assessment of parameter stability and predictive robustness by refitting the model on nearly the full dataset while testing its generalization on the held-out observation. The refitted parameters for each LOO iteration $i$ are denoted as $\hat{\theta}^{(i)}$. For each candidate parameter set $\hat{\theta}^{(i)}$, we compute the likelihood using all observations $y$ to ensure that the sample size is consistent with the null model in the downstream calculation of Bayesian information criterion (BIC) (13):

$$\mathcal{L}^{(i)} = P(y \mid \hat{\theta}^{(i)}) \tag{23}$$

Given that each refitted parameter set is assigned equal weight $w_i = \frac{1}{n}$, the log likelihood (14) combining all LOO models is calculated as:

$$\log P(y \mid \hat{\theta}) = \log\left(\sum_{i=1}^{n} w_i \mathcal{L}^{(i)}\right) = \log\left(\frac{1}{n}\sum_{i=1}^{n} \mathcal{L}^{(i)}\right) \tag{24}$$
